# A comparative assessment of aging-related NADPH diaphorase positivity in the spinal cord and medullary oblongata between pigeon and murine

**DOI:** 10.1101/650457

**Authors:** Yunge Jia, Wei Hou, Yinhua Li, Xiaoxin Wen, Chenxu Rao, Zichun Wei, Tianyi Zhang, Xinghang Wang, Xiuyuan Li, Lu bai, Weijin Zhang, Pan Wang, Jing Bi, Anchen Guo, Jie Wang, Huibing Tan

## Abstract

NADPH diaphorase (N-d) positive neurons has been examined in many animals. N-d neurodegenerative neurites were detected in some animal models. However, detailed information of N-d positivity and aging related changes was still lack in the spinal cord and medulla oblongata of pigeons. In this study, we evaluated the N-d positivity and aging alterations in the spinal cord and medullary oblongata of the pigeon compared with rat and mouse. In pigeons, N-d neurons were more numerous in the dorsal horn, around the central canal and in the column of Terni in the thoracic and lumbar segments and scattered neurons occurred in the ventral horn of spinal segments. N-d neurons also occurred in the white matter of spinal cord. Morphometrical analysis demonstrated in the lumbosacral, cervical and thoracic regions. Compared with young pigeons, the size of N-d soma was significantly altered in aged pigeons. Meanwhile, the dramatic morphological changes occurred in the lumbar to sacral segments. The most important findings of this study were aging-related N-d positive bodies (ANB) in aged pigeons, mainly in the nucleus cuneatus externus (CuE), occasionally in the nuclei gracilis et cuneatus. ANBs were identified in the gracile nuclei in spinal cord in the aged rats and mice. ANBs were also detected in the CuE spinal nucleus in the aged rats. Immunohistochemistry also showed that the aging changes occurred in the cell types and neuropeptides in aged animals. The results suggested the weak inflammation and neuronal dysfunction in the spinal cord in aged pigeons. Our results suggested that the ANB could be considered as aging marker in the central nervous system.

## INTRODUCTION

Nicotinamide adenine dinucleotide phosphate diaphorase (N-d) positive neurons are distributed in the spinal cord, besides the central nervous system (Valtschanoff et al., 1992; Vizzard et al., 1994a; Crowe et al., 1995; Marsala et al., 1998; Marsala et al., 1999). N-d activity has also been considered identical to nitric oxide synthase (NOS) in neurons (Dawson et al., 1991; Hope et al., 1991; Young et al., 1997). Indeed, N-d histochemistry correlates with NOS immunocytochemistry (Bruning, 1993; Moreno et al., 2002). However, some of the experiments reveal that NOS is not always matched with N-d histochemistry (Traub et al., 1994; Vizzard et al., 1994b; Pullen and Humphreys, 1995; Tan et al., 2006). N-d does not co-localize with NOS in nearly half of cat sacral motoneurones (Pullen and Humphreys, 1995). Nevertheless, the N-d histology is used to identify and describe the distribution of putative neuronchemistry (Scherer-Singler et al., 1983; Rice, 1995).

In the avian brains, N-d histochemistry has been considerd to map NOS-positive neurons (Bruning, 1993; Ambalavanar et al., 1994; Cozzi et al., 1997). The distribution of N-d pigeon neurons is studied in the brain and the cervical segment (Atoji et al., 2001a). The other segments of the spinal cord are still important. The avian spinal cord shows some particularity in the lumbosacral region that discloses dorsally to form a rhomboid sinus occupied by the glycogen body (Matulionis, 1972; Necker, 1999; Moller and Kummer, 2003; Imagawa et al., 2006). Specialized semicircular canal-like structures (lumbosacral canals) is noted the coronal sections at the rhomboid sinus (Necker, 1999). Furthermore, many paragriseal neurons occur in the white matter of lumbosacral spinal cord. In rodents, N-d neurons in the caudal spinal cord is important to the regulation of pelvic organs. Different to other mammal species, pigeon peripheral organs of lumbosacral spinal cord develop cloaca which also innervates with N-d neurons (Atoji et al., 2001b). Several studies reveal N-d neurons in the lamina II of the dorsal horn in the pigeon (Necker, 2004). The cells of origin of sympathetic preganglionic fibers are localized to a well-defined nucleus dorsal to the central canal (column of Terni) (CT) (Macdonald and Cohen, 1970). In general, parasympathetic preganglionic nuclei locate in the lumbosacral segments caudally to sympathetic nuclei. Necker demonstrates that N-d positivity in the spinal cord of the pigeon is comparable to that in mammalian species despite divergence (Necker, 2004). Actually, it is indicated that there is an anatomical discrepancy of the dorsal commissural nucleus (DCN) in the rodent lumbosacral spinal cord (Tan et al., 2006). The similar anatomical position of the dorsal commissure of lumbosacral segments in pigeons occurs an open specialization of the central canal as demonstrated above. In mammal species, N-d neurons in the DCN correlate with the regulation of the pelvic organs (Vizzard et al., 1994a; Nadelhaft and Vera, 1996). Injury or dysfunction of urogenital visceral organs may be involved in the lumbosacral spinal cord (Birder et al., 1999; Fowler et al., 2008).

The significance of N-d reaction is used to characterize some neuropathology in aging deterioration and neurodegenerative disease (Yamada et al., 1996; Kuo et al., 1997; Reuss et al., 2000; Quinn et al., 2001; Tan et al., 2006). In the cerebral cortex (Yamada et al., 1996) and striatum (Yamada et al., 1996; Kuo et al., 1997), age-related changes of N-d-positive neurons occur in aged rats with no significant changes in the hippocampus. Aging related changes of N-d histochemistry demonstrate to be regionally specific in the rat (Mizukawa et al., 1988; Kuo et al., 1997). Our previous study shows that N-d histochemistry of aging alterations demonstrate a specific regional changes in the lumbosacral spinal cord of aging rats, in which the N-d spheroids were termed as “aged-related N-d positive bodies (ANB)” (Tan et al., 2006). Recently, we found that specialized meganeurites or megaloneurites which were different from the alteration of the aged rats occurred in the sacral spinal cords in aged dog (Li et al., 2018). Birds are considered as an aging model in biomedical research (Coppola et al., 2015), aging-related changes of brain and sensory mechanisms has been studied in homing pigeons (Meskenaite et al., 2016). However, to our knowledge, little attention has been paid to the N-d reactivity in medulla oblongata of the aged pigeon and in all the spinal segments of the pigeon with reference to the lumbosacral segment and the difference between the young and aged pigeons (Atoji et al., 2001a; Necker, 2004). The aim of the present study was to investigate the cellular types and quantification of N-d-containing neurons in all regions of the spinal cord and morphological aging changes from the lumbar to the sacral segment. A relative semi-quantification is used to explain the significance of their distributions. The most important thing was that to explore whether there was age-related NADPH-d positive body (ANB) in the lumbosacral spinal cord、GR、 CU of aged pigeons and mice. Was ANB as a neurodegenerative marker in different species? By comparing the similarities and differences among aged pigeons, aged rats and aged mice, the far-reaching significance of the existence of ANB was further elaborated.

## MATERIALS AND METHODS

### Animals and Tissue Preparation

Experiments were performed using aged 9 – (n = 5), 14 – (n = 5), 16 – (n = 5), young 1 – (n = 5) year of the racing pigeons (Columba Livia) and aged 18 – (n = 10), young 2 – (n = 6) month of the rats (Sprague Dawley) and aged 18 – (n = 6), young 2 – (n = 6) month of the mice (C57BL/6J). Every pigeon had identified number of leg ring, which was used for age confirmation. The pigeon fanciers had approved certification for breeding the pigeons and taken part in pigeon racings as club member. Pigeons were housed in farming facilities. Food and water were provided properly. All murine were provided by the Animal Laboratory Center of Jinzhou Medical University. All experimental procedures were approved by the Ethics Committee in Animal and Human Experimentation of the Jinzhou Medical University.

For physical examination of pigeons, weight loading was determined to stop flight and reduced the loading weight until the pigeon restored ability to flight. Behavioral tests were also examined with hot plate test and von Frey before sacrifice of all of the pigeons. For the thermal test of responses to the nociceptive heat stimuli, the graded temperature of the device (Zhenghua, China) was set from 52-60°C. For test of a mechanical response, von Frey hairs (Yuyanyiqi Touch test, China) were selected to test withdrawal paws of pigeons in a chamber.

All animals were anesthetized with pentobarbital sodium (45 mg/kg, i. p.). The chest cavity was opened and a cannula was inserted into the ascending aorta via the left ventricle. Perfusion was performed using 100–150 ml of 0.9% NaCl followed by 4% paraformaldehyde in 0.1M sodium phosphate buffer (PB, pH 7.4). The spinal cord and brain were rapidly removed and postfixed with 4% paraformaldehyde in 0.1 M phosphate buffer, left at 4°C for 2h and then placed in 30% sucrose overnight.

### Histochemical processing

Staining was performed using free-floating sections. The spinal cord from the cervical to coccygeal segments and medulla oblongata were cut coronally into 40µm thick sections using a Leica CM1950 cryostat. Part of the slice was stained and examined by N-d histochemistry. For N-d histochemistry, free-floating sections were incubated in PB containing Triton X-100 (0.3%), NADPH (1 mg/ml, Sigma) and nitroblue tetrazolium (1 mg/ml, Sigma) at 37°C for 90 to 120 min. The reaction was stopped by washing the sections with the phosphate buffered saline (PBS, 0.01M). The sections were examined and photographed under an optical microscope (Olympus BX35, camera DP80, Japan).

For Nissl staining, the frozen sections were stained with 0.1% cresyl violet for 10-20 min, rinsed with PBS, dehydrated in a graded alcohol series, cleared with xylene and mounted with neutral gum. The sections were observed using light microscopy.

### Immunofluorescence staining

For immunofluorescence staining, sections were incubated in 3% normal goat serum in PBS for 1h and incubated overnight at 4°C with the following primary antibodies: (1) rabbit monoclonal antibody against NeuN (1:1000, MAB377, Millipore), (2) mouse anti-GFAP (1:1000, SAB5201114, Sigma) for glial fibrillary acidic protein, (3) mouse anti-Iba1 (1:800, 1919741, wako) for microglia/macrophages, (4) rabbit anti-VIP (1:1000, HPA017324, Sigma) for vasoactive intestinal peptide, (5) mouse anti-MAP2 (1:200, NB600-1372, NOVUS) for microtubule-associated protein 2, (6) mouse anti-calcitonin gene-related peptide (CGRP; 1:100, Sigma, USA) of neuronal peptide in the nervous tissues. In each immunofluorescence testing, a few of sections were incubated without primary antibody as a negative control. After extensive washing, sections were incubated with Alexa Flour 594 anti-rabbit IgG (1:800, A11037, Life) and or Alexa Flour 488 anti-mouse IgG (1:800, A11001, Life) at room temperature for 1h and mounted with DAPI (F6057, Sigma). The sections were observed using fluorescence microscope (Olympus BX35, camera DP80, Japan).

### Qualitative and quantitative analyses and stereological measurement in pigeons

The spatial distribution of N-d neurons in corresponding sections was obtained from cervical, thoracic, lumbar and sacral segments and medulla oblongata from each pigeon. Within each spinal segment, 30 sections were randomly selected for analysis. The number of cells was counted in the following regions: dorsal horn (lamina □-□), central canal (lamina □ and □) and ventral horn (lamina □ and □). Within each medulla oblongata segment, 10 sections were randomly selected for analysis. The number of cells was counted in the CuE.

For stereological measurement, a random section sampling of N-d staining was performed under the microscope (Leica DM6B) with motorized 3-axis stage control by the software (Stereo-Investigator MBF Bioscience, Williston, VT). A researcher blinded to the animals traced a contour around the location for measurement under 10X objective (N.A=0.30) and counting under 40X objective (N.A=0.75). The cells were counted when nucleus firstly came into focus inside the counting frame. Neuronal numbers and areas of cells were measured for the quantitative analysis. The differences among groups were statisticed by one-way analysis of variance (ANOVA) followed by the Newman-Keuls post hoc test. The statistical significance threshold was set at P < 0.05. The following parameters, which included cell body area (values expressed in µm^2^). Mean values were expressed as mean ± SEM.

### Measurement for immunofluoresence in pigeons

Semi-quantitative analysis with ImageJ software (version 1.48). Sections (20 young and 20 aged) of lumbosacral spinal cord were stained with NeuN, GFAP, Iba1, MAP2, VIP. We selected the area of 150μm × 200 μm from the images and calculated the percentage of the total area of all fluorescent reactions in the area. Percentage multiplied by 100 as statistical data. Data was compared using two-tailed unpaired T test, using GraphPad Prism software. A P-value less than 0.05 was considered statistically significant.

## RESULTS

To assess pain sensitivity by the hot plate test, there was no significant effect of age on paw responses to a thermal stimulus with the upper cut-off at 60°C over 30 seconds. This result indicated that pigeons were insensitive to thermal nociception and no age-related changes in pain sensitivity. Aged pigeons also had no significant change of threshold to von Frey hairs compared to young rats selected 300g-hair as the upper cut-off for testing. Suggested different to rodents, ordinary behavioral tests do not suitable for pigeons to test mechanothermal sensation. One fourth body weight was loaded on the pigeons to determine the ability to flight and revealed that there were no significant differences between the young and aged pigeons. There is also no significant change in neuromuscular strength in body weight loading experiment.

N-d positive neurons were centrally distributed in the dorsal horn, CT, near central canal, scattered in the ventral horn, white matter and the Clarke’s column (CIC) of enlargement (Supplementary Figure S1A). The N-d positivity of the Lissauer’s tract and the lateral collateral pathway were not clearly detected in both young and age pigeons in our experiments, which is well defined in the cats (Vizzard et al., 1994b). N-d staining bands of dense N-d-positive fibers were identified in the dorsal horn of all spinal cord segments. It was worth noting that the avian preganglionic CT was located dorsal to the central canal and was present in the thoracic segments. Coincidentally, there was a similar group of cells in the corresponding position of the lumbar segments (Supplementary Figure S1B).

**Figure S1.**
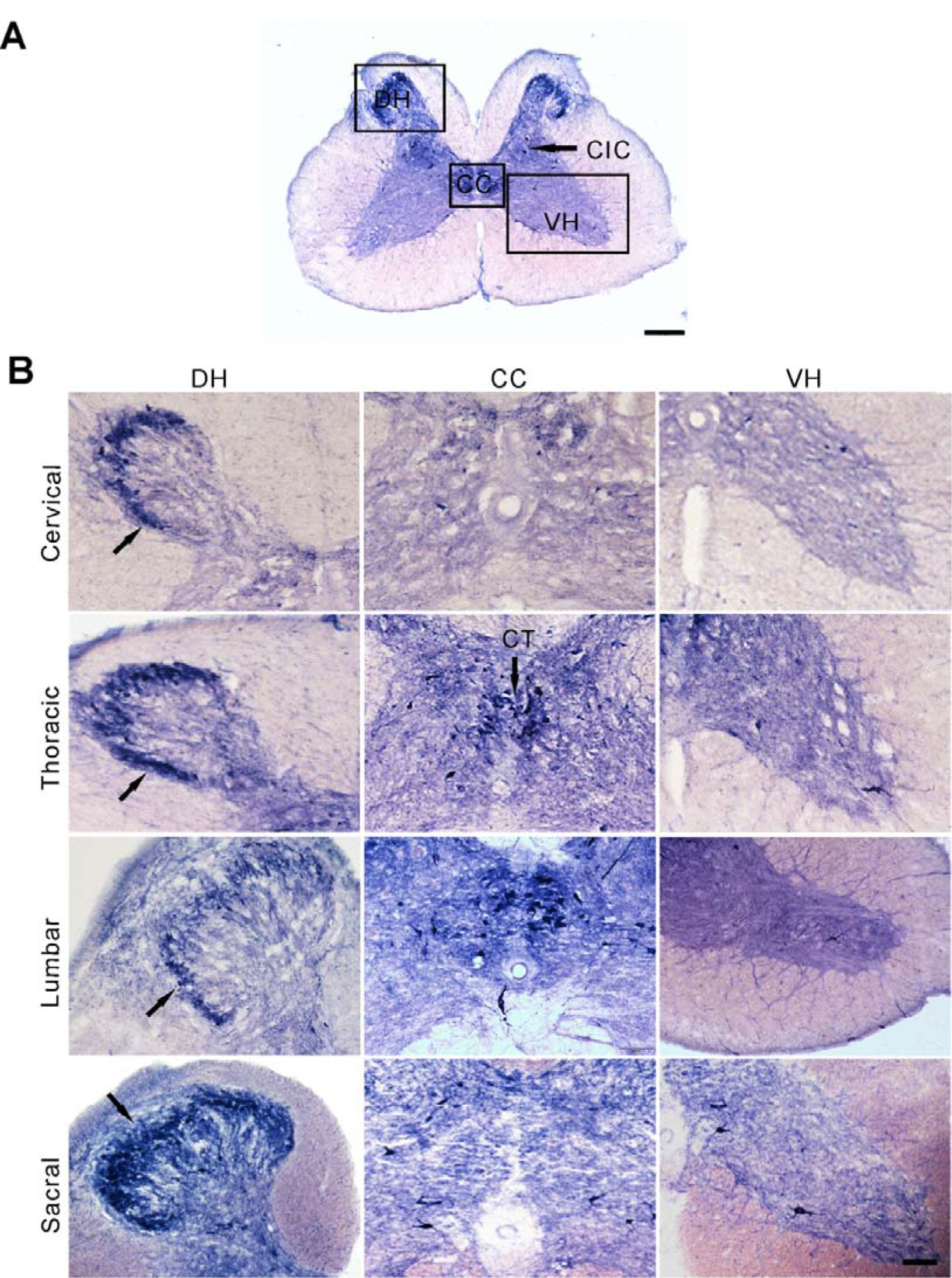
Regional indication for example for N-d staining of the pigeon spinal cord. (A) Square boxes indicate sample areas in the lumbar segment of young adult pigeons. (B) N-d regional distribution in different spinal segments. Black arrows indicate densely-stained band in the dorsal horn. DH: dorsal horn, CC: central canal, VH: ventral horn, CT: column of Terni, CIC: Clarke’s column. Scale bar, 200μm for A and 50μm for B.

In general, N-d histochemistry produces Golgi-like staining (Valtschanoff et al., 1992). According to the number of dendrites, two subgroups (types) were identified. Type □ was stained strong intensity with massive formazan deposits by N-d staining throughout the cytoplasm and the nucleus and detected more than one dendrite. Type □, on the other hand, had less than 2 or no dendrites (Figure 1A). Type □ accounted for ninety percent of the number of N-d neurons. Most of the N-d cells were located in the lamina □, while a very small number of neurons were detected in lamina □ and lamina □ (Figure 1B). The CIC was obviously detected in lamina □ in the lumbosacral enlargement. The N-d positive cells in this region were large in size and stained few dendrites. In addition, the number of cells in this region was 2-5 per section (Figure 1C). There was a group of N-d neurons between the dorsal column and central canal in the thoracic and lumbar segments, most of which were type II cells (Figure 1D, E). The number of neurons in the white matter was counted from 30 slices in each spinal segment (Figure 1F, G). The number of neurons in the white matter was increased from the cervical to the sacral segments (Figure 1H). The N-d staining of dorsal root ganglia (DRG) was relatively uniform. Majorities of the cells were positively stained in the DRG (Figure 1I).

**Figure 1.**
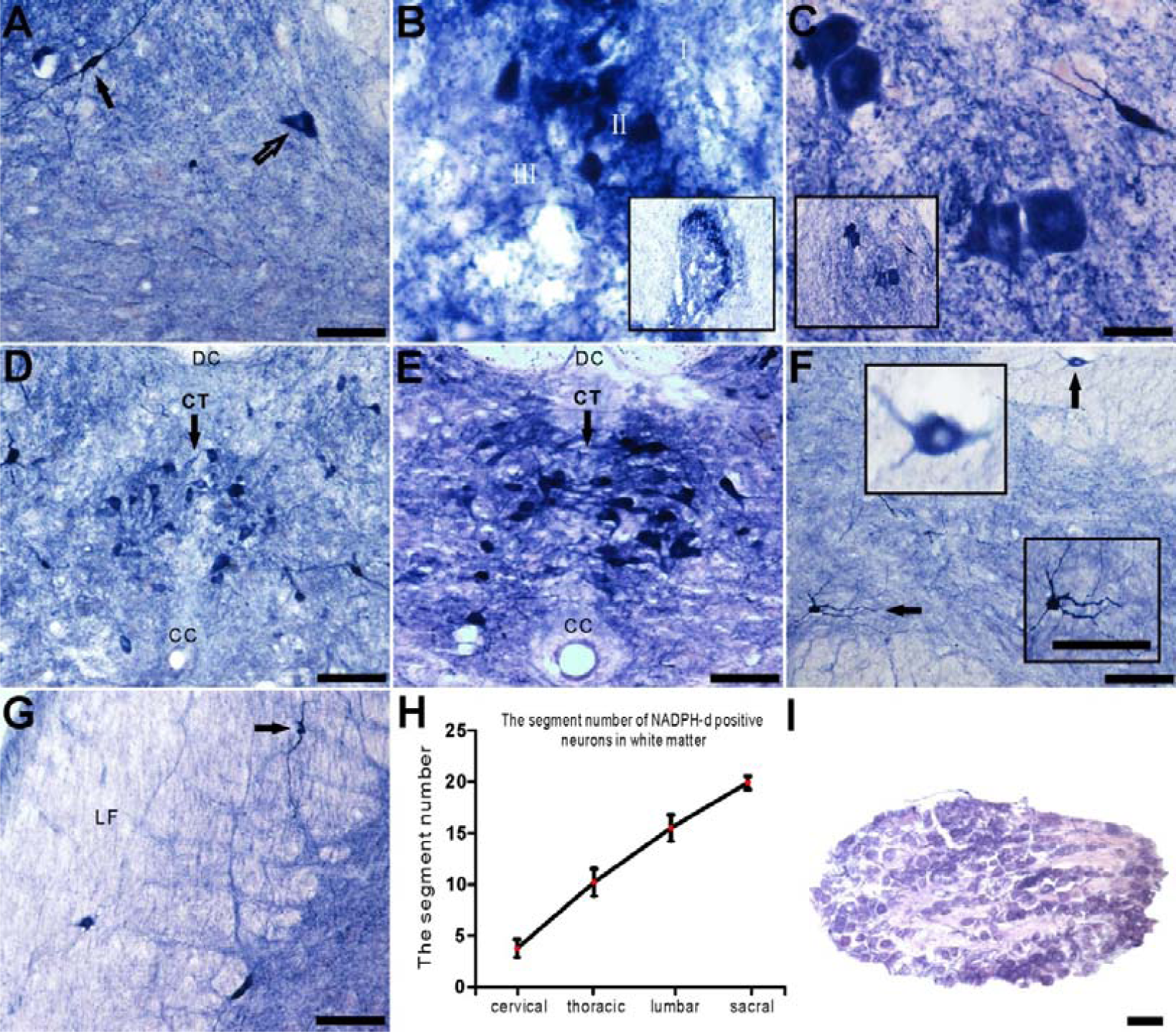
N-d positive neurons in the spinal cord of young adult pigeons. (A) Classification of N-d neurons. The black arrow indicates Type I neuron while open arrow indicates type II neuron. (B) Dyed bands of the dorsal horn in the cervical spinal cord. (C) Clarke’s column (CIC) in the lumbar. (D) Positive cells in the column of Terni (CT) in thoracic segment. (E) Magnification of the intermediate region in lumbar segment, a group of labeled neurons in the similar region named Terni (CT) in thoracic segment. The arrows embedded in the same graph indicated an area to be magnified F. (F, G) N-d neurons in the ventral horn and the white matter of the lumbar segment. (H) The number of neurons in the white matter for each spinal segment. (I) The DRG cells of lumbar segment. DC: dorsal column, CC: central canal. Scale bar=50μm.

Although the N-d cells were detected in both gray matter and white matter of the spinal cord, it was not evenly distributed (Figure 2A). In a certain segment, the number of N-d positive cells in the dorsal horn was greater than in that around the central canal and ventral horn respectively (*p* < 0.01). In addition, comparing the regional distribution, the number of N-d cells in the central canal was greater than that in the ventral horn, but in the cervical segment, the cellular distribution was opposite (*p* < 0.05, Figure 2B). The number of N-d cells also varied in distinct regions of all spinal segments. For the dorsal horn, the largest number of N-d positive cells was especially found in the lumbar and sacral segments. Around the central canal, the number of cells in the thoracic and lumbar segments was more than that in the cervical and sacral segments. For the ventral horn, the number of cells in the thoracic and cervical segments was more than that of the lumbosacral segment (*p* < 0.01, Figure 2D). Next, we analyzed morphological criteria of perikaryon area. The area of positive cell body in the ventral horn was the largest in all spinal segments followed by the central canal and the smallest area of cell body was that of dorsal horn (*p* < 0.05, Figure 2C). For the same anatomical regions such as the dorsal horn of the different segments, the perikaryon area was also diverse. The cell body area in the lumbosacral segments was larger than that in the dorsal horn, and similar results also revealed in the ventral horn and in the central canal (*p* < 0.05, Figure 2E).

**Figure 2.**
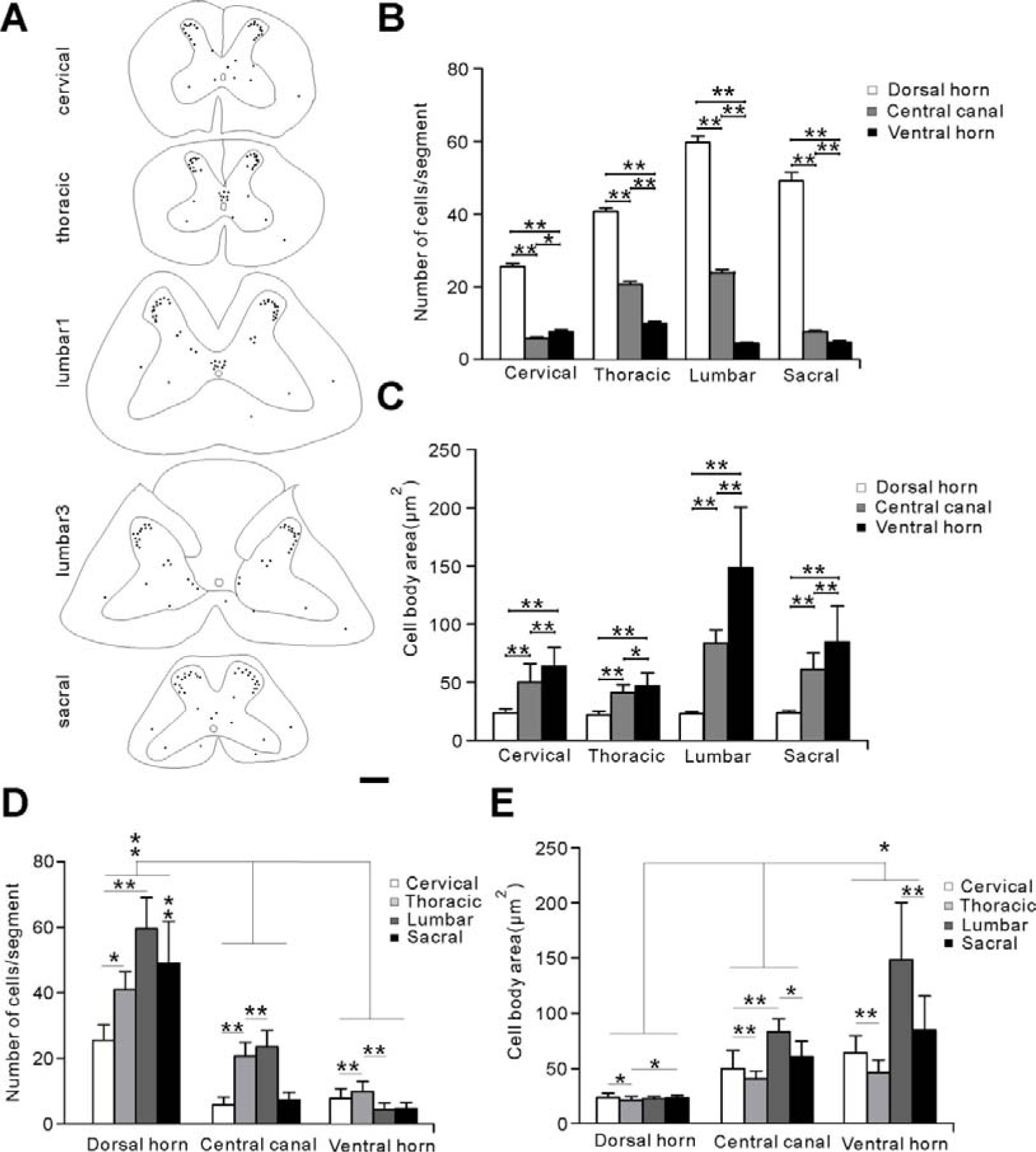
Morphological creteria of N-d neurons of young adult pigeons. (A) Pattern of the segmental distribution of N-d neurons. Dots indicate neuronal soma. (B, C) Statistical analysis of cell number and cell body area in each segment. (D, E) Statistical analysis in the same parts of different segments of the spinal cord. (\**p* < 0.05; \*\**p* < 0.01). Scale bar=200μm.

The lumbosacral enlargement includes the very obvious rhomboid or lumbosacral sinus as a specificity of all birds, which enlarged the spinal nervous tissue because of glycogen body enclosed in a dorsal open space (rhomboid sinus). From the lumbar to the sacral segment, the morphological change was splitting of the dorsal spinal parenchyma from opening to closure indicated with N-d staining (Figure 3). As mentioned above, the specialization of pigeon spinal cord was a semicircular canal-like structure which was occupied by the glycogen body in the dorsal lumbosacral segments. At the rostral lumbar segment, the N-d positive cells between the dorsal column and central canal occurred in cluster arrangement (Figure 3B). Compared with Figure 3B, there was a change above the central canal and a lightly stained band vertically oriented (Figure 3C-E). The lightly stained band was further widened, a splitting space appeared, and was filled with colloidal substance (Figure 3E) until the two sides of the gray matter and the dorsal column were completely separated (Figure 3F). The number of type II neurons decreased until it eventually disappeared. Meanwhile, the small amount of type I cells that scattered did not apparently change. After the combinative component of dorsal gray matter transitionally disappeared, the parenchyma cells on both sides were stilled connected by neurons in the anterior gray matter, as shown in Figure 3I. The dorsal part of lumbosacral spinal cord forms a rhomboid sinus which set in the glycogen body. When it caudally merged to the closure of rhomboid sinus, the commissural positive fibers were detected in the gray matter on both sides of the gray matter (Figure 3G). Finally, the two sides of the separated dorsal columns were fused. Thus, the rhombus sinus was completely closed again (Figure 3H).

**Figure 3.**
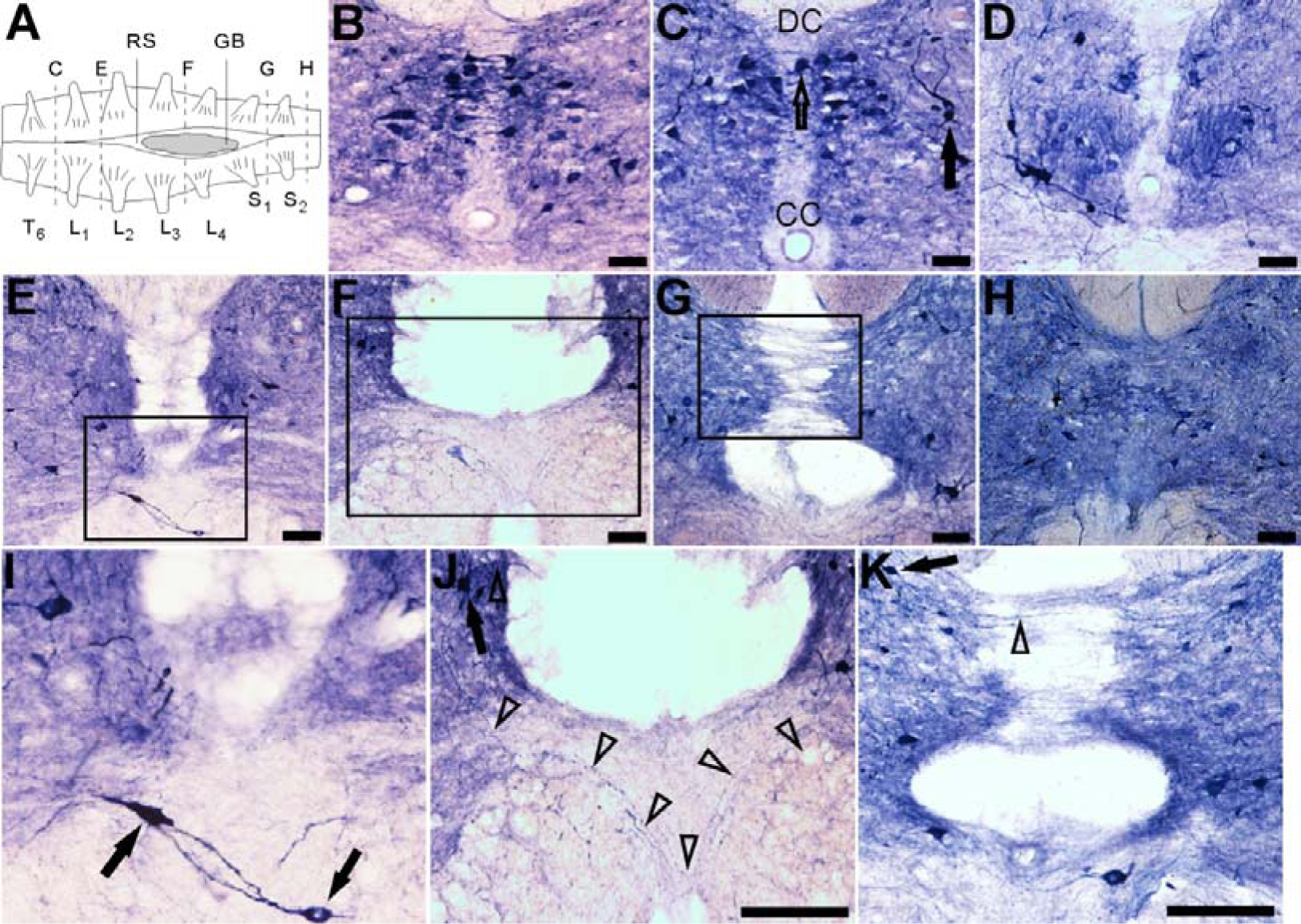
Morphological profile of the central canal, anterior and posterior gray commissure and dorsal column in the lumbosacral spinal cord of young adult pigeons. (A) Illustration shows the dorsal view of the lumbar enlargement of the spinal cord. The dotted lines are the corresponding sections. Coronal sections are indicated from rostral to caudal orientation (B-H). (I-K) Shows a higher magnification from boxes in E-G. The black arrow indicates type I N-d neurons and the open arrow shows type II in C. (I) The black arrow shows positive neurons located in the anterior commissure. (J) Neurons in intermediomedial region (black arrow) which sent processes (open arrowheads) into the anterior commissure and cross to the opposite side. (K) Labelled neurons in intermediomedial region (black arrow) which sent fibers into the rhomboid sinus (open arrowheads). CC: central canal, DC: dorsal column, RS: rhomboid sinus, GB: glycogen body. Scale bar=50μm.

N-d histochemical and Nissl staining revealed with related cytoarchitecture in the GC and CuE by studying the medulla oblongata of young and aged pigeons (Figure 4A-C). Normal N-d positive cells were found in the CuE of young pigeon (Figure 4D). Furthermore, there were N-d positive neurons with Golgi-like dendrites (Figure 4D1). Importantly, there were N-d positive bodies in the CuE of aged pigeon. The N-d spheroids were termed as “aged-related N-d positive bodies (ANB)”. The number of ANBs was increased with age (Figure 4E-G). The ANBs were characterized with diverse shapes (Figure 4E1-G3).

**Figure 4.**
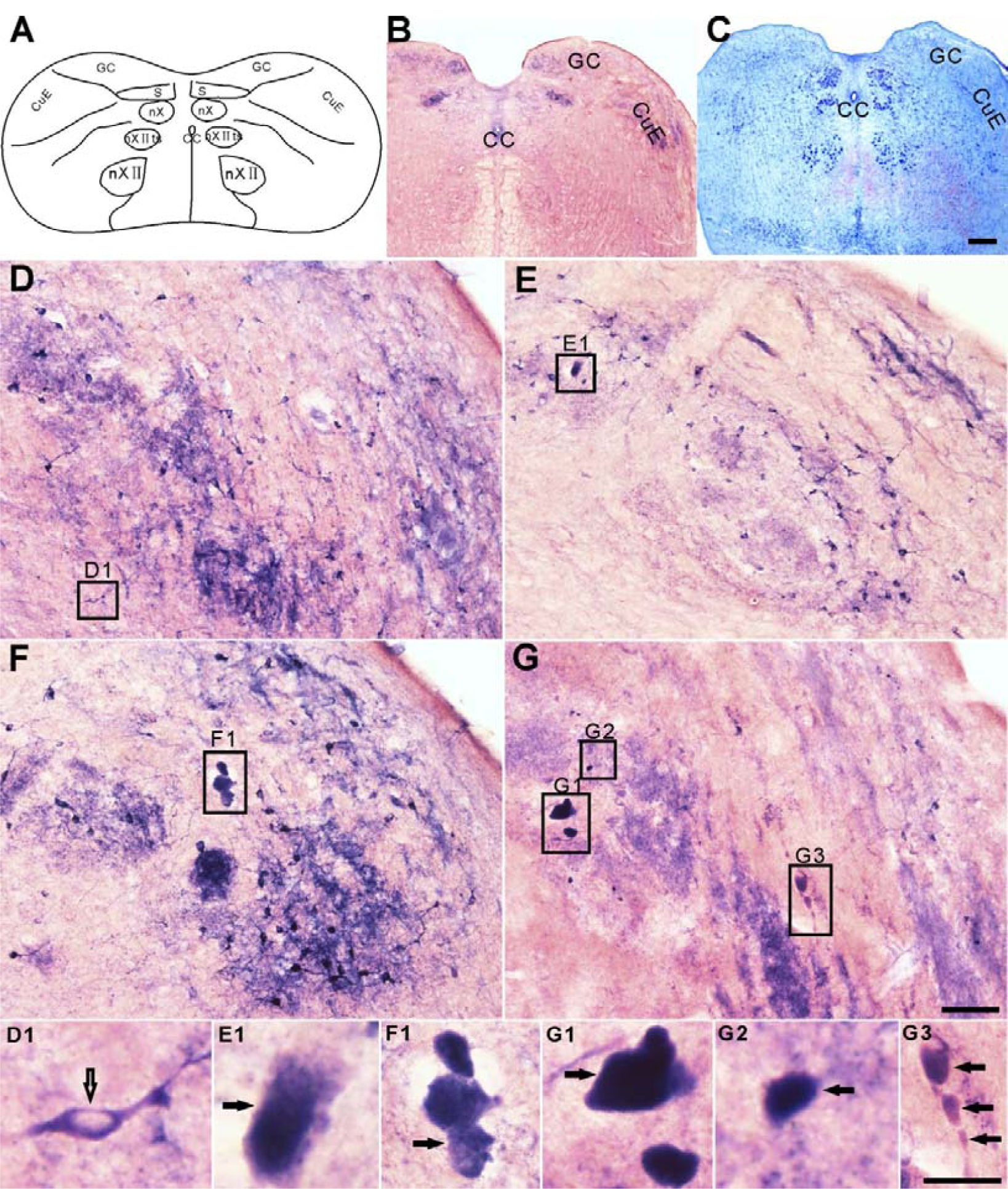
Swelling N-d neurites detected in the medulla oblongata of the aged pigeons. (A) Schematic illustration of the transverse plane through gracile nuclei (GC) and external cuneate nucleus (CuE) of the medulla oblongata level. (B) N-d reactivity in medulla oblongata level of young pigeons. (C) Nissl staining of medulla oblongata at the same level of young pigeons. (D) N-d reactivity in CuE of young pigeons (1-year old). (E) N-d reactivity in CuE of aged pigeons (9 years old). (F) N-d reactivity in CuE of aged pigeons (14-years old). (G) N-d reactivity in CuE of aged pigeons (16-years old). (D1) Normal N-d positive neurons in young pigeons (open arrow indicates). Higher magnifications of ANB in CuE (E1-G3) with various shapes (black arrow shows). Abbreviations: GC, nuclei gracilis et cuneatus; S, nucleus tractus solitaries; CuE, nucleus cuneatus externus; nX, nucleus motorius dorsalis nervi vagi; nXIIts, nucleus nervi tracheaosyringealis; nXII, nucleus nervi hypoglossi; CC, central canal. Scale bar 200μm for B and C; 50μm for D-G; 10μm for D1-G3.

Although ANB was found in the CuE, but there was still the existence of normal N-d positive neurons of aged pigeon. The number of N-d positive neurons was decreased in the CuE with age (Figure 5A), while the number of ANB was increased with age (Figure 5B).

**Figure 5.**
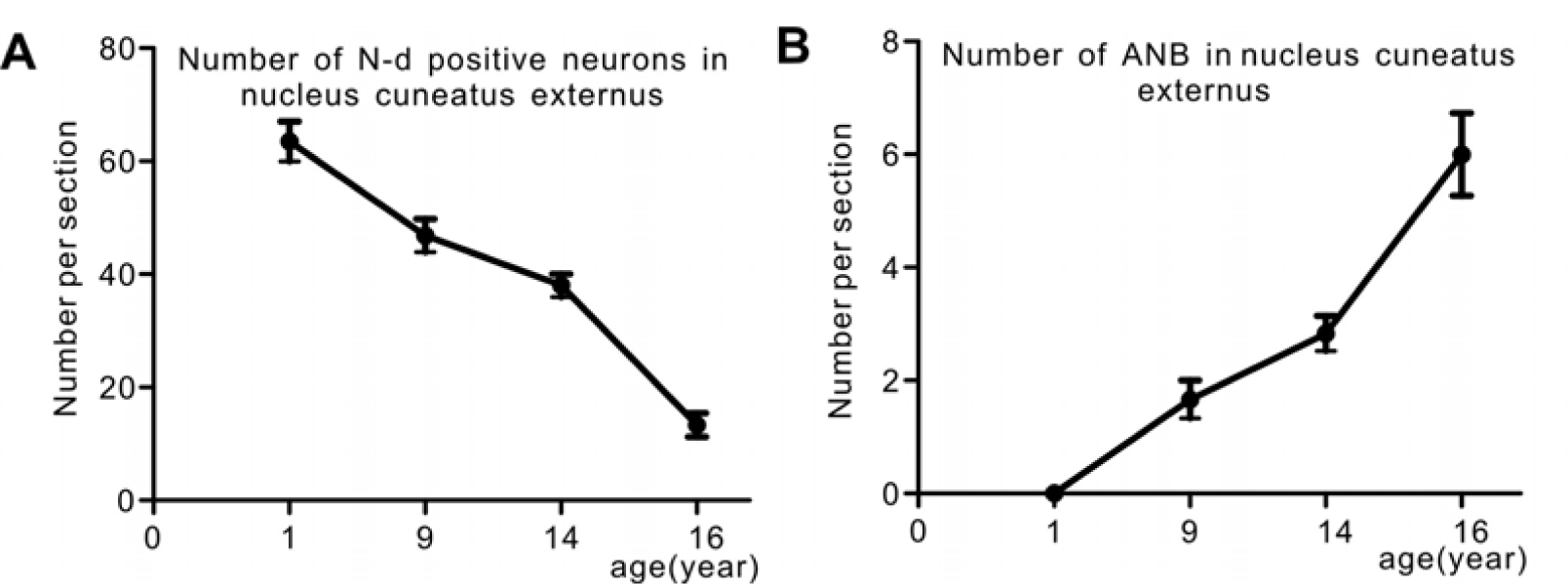
Histograms show the number of N-d positive neurons and ANB in nucleus cuneatus externus with age. (A) the number of N-d positive neurons in nucleus cuneatus externus was decreased with age. (B) the number of ANB in nucleus cuneatus externus was increased with age.

Although birds and mammals were two very different species, but ANB has been found in the medulla oblongata of both two species (Figure 6). It was worth mentioning that the ANB in pigeons mainly distributed in the CuE and occasionally in the GC (Figure 6A, C), which indicated dramatically in contrast to the ANB in rats. There were numerous ANB mainly distributed in the gracile nucleus (GR) (Figure 6B, D) and a small amount in the cuneatus nucleus (CU) as well as CuE (data not showed here).

**Figure 6.**
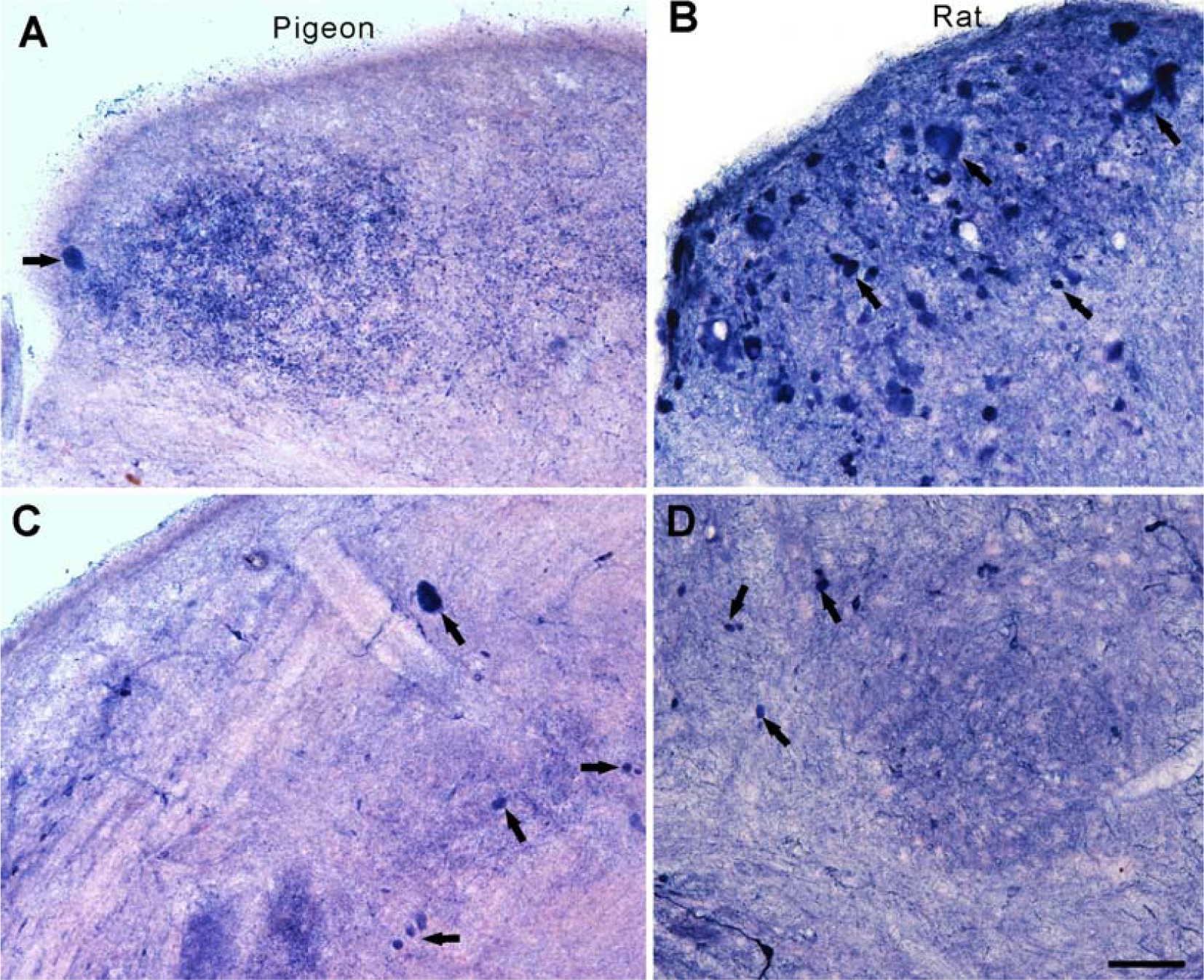
ANB in the dorsal column of aged pigeons and aged rat. (A) Nucleus gracilis of aged pigeons. (B) Nuclei gracilis of aged rat. (C) Nucleus cuneatus externus of aged pigeons. (D) Nucleus cuneatus of aged rat. Black arrow indicates ANB. Scale bar=50μm.

We used N-d staining to assess the difference of spinal cord between young and aged pigeons (Figure 7). According to our previous studies in aged rats (Tan et al., 2006) and dogs (Li et al., 2018), no aging-related N-d degenerative neurites were found in the present study in the lumbosacral segment of aged pigeons (Figure 7A). Consistent with previous studies (Tan et al., 2006). ANB was also found in the lumbosacral segment of aged rats (Figure 7B). Morphological measurement in the CT and CIC was taken for the number of neurons and the cell body area (Figure 7C-G). Results showed that the number of cells in the CIC between young and aged was unchanged (*p* > 0.05), but the cell body areas in the column of the CT and CIC, and the number of cells in the CT had significantly decreased in that of aged pigeons (*p* < 0.05). Compared with the number of neurons in dorsal horn between the young and aged spinal cord, it was found that except for the thoracic segment, the number of cells in the other segments decreased in that of aged pigeons (*p* < 0.01, Figure 7G).

**Figure 7.**
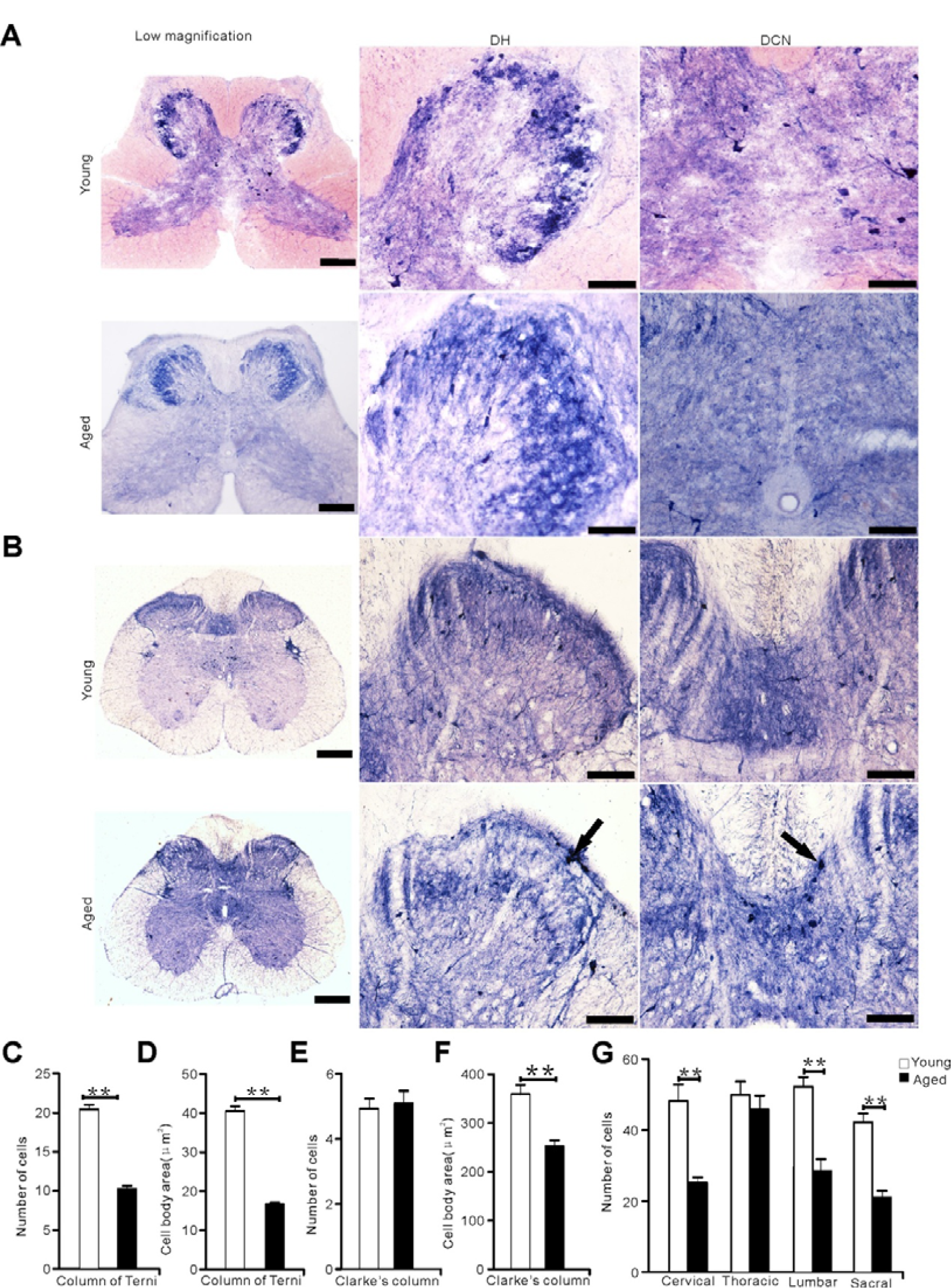
N-d reactivity in the sacral segment between young and aged animals (pigeons and rats). (A) N-d positive cells in the sacral segment of young and aged pigeons. (B) N-d reactivity in the sacral segment of young and aged rats. Black arrow indicates ANB. (C-G) analysis of N-d positive cells in the dorsal horn, column of Terni (CT), Clarke’s column (CIC) of the spinal cord between young and aged pigeons. (C) Number of N-d positive cells in the column of Terni (CT). (D) Cell body area in the column of Terni (CT). (E) Number of N-d positive cells in Clarke’s column. (F) Cell body area in Clarke’s column. (G) Number of N-d positive neurons in dorsal horn of spinal cord between young and aged pigeons. DH: dorsal horn, DCN: dorsal commissure nucleus. \**p* < 0.05, \*\**p* < 0.01. Scale bar=50μm.

The sensory fibers from DRG terminate in GR, CU or CuE through gracile fasciculus and cuneate fasciculus, while ANB was found in GR, CU and CuE of aged rats and pigeons. The ANBs were sporadically detected in the dorsal column of each spinal cord segment in the aged pigeons (Figure 8A, B). Consequently, the ANBs occurred in GR, CU and CuE of the aged pigeons. No ANB was detected in the dorsal column of young pigeon (Figure 8C). Comparative N-d histochemical staining of aged rats, ANBs were detected in the cervical, thoracic segment (Figure 8D, E).

**Figure 8.**
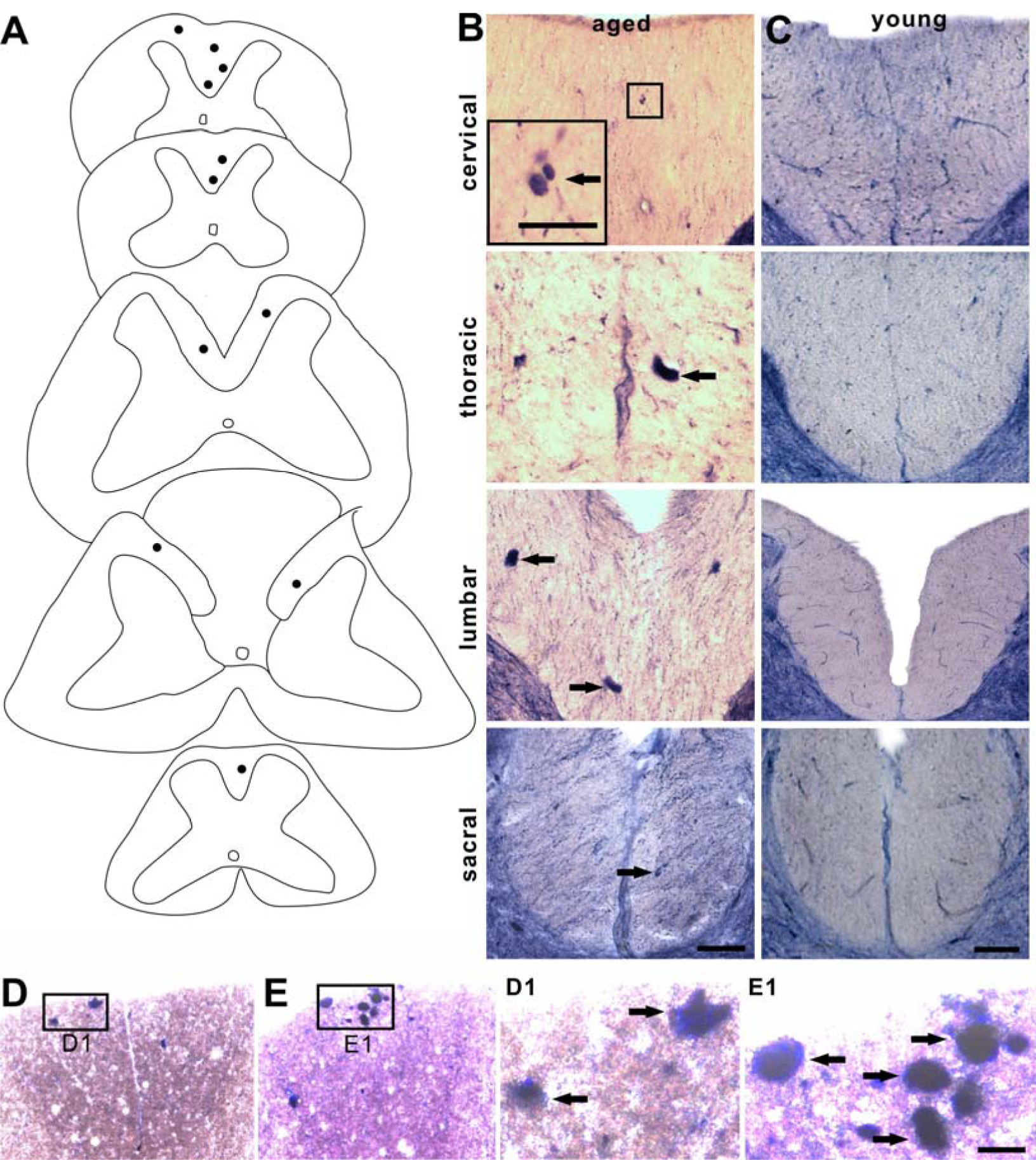
Neurodegenerative N-d reactivity in dorsal column compared between the aged pigeons and rats. Distribution of ANBs at cervical, thoracic, lumbar, sacral segment of aged pigeons (A). Dots indicate ANBs in A. The photo images of dorsal column in aged and young (C) pigeons. ANB in the dorsal column of cervical (D, D1) and thoracic (E, E1) segment of aged rats. Black arrow was referred to ANB. Sacral bar =50μm.

Next, ANBs were confirmedly detected found in GR and DCN regions of lumbosacral spinal cord in aged mice (Figure 9). This indicated the significance of ANB’s ubiquitous existence in different species.

**Figure 9.**
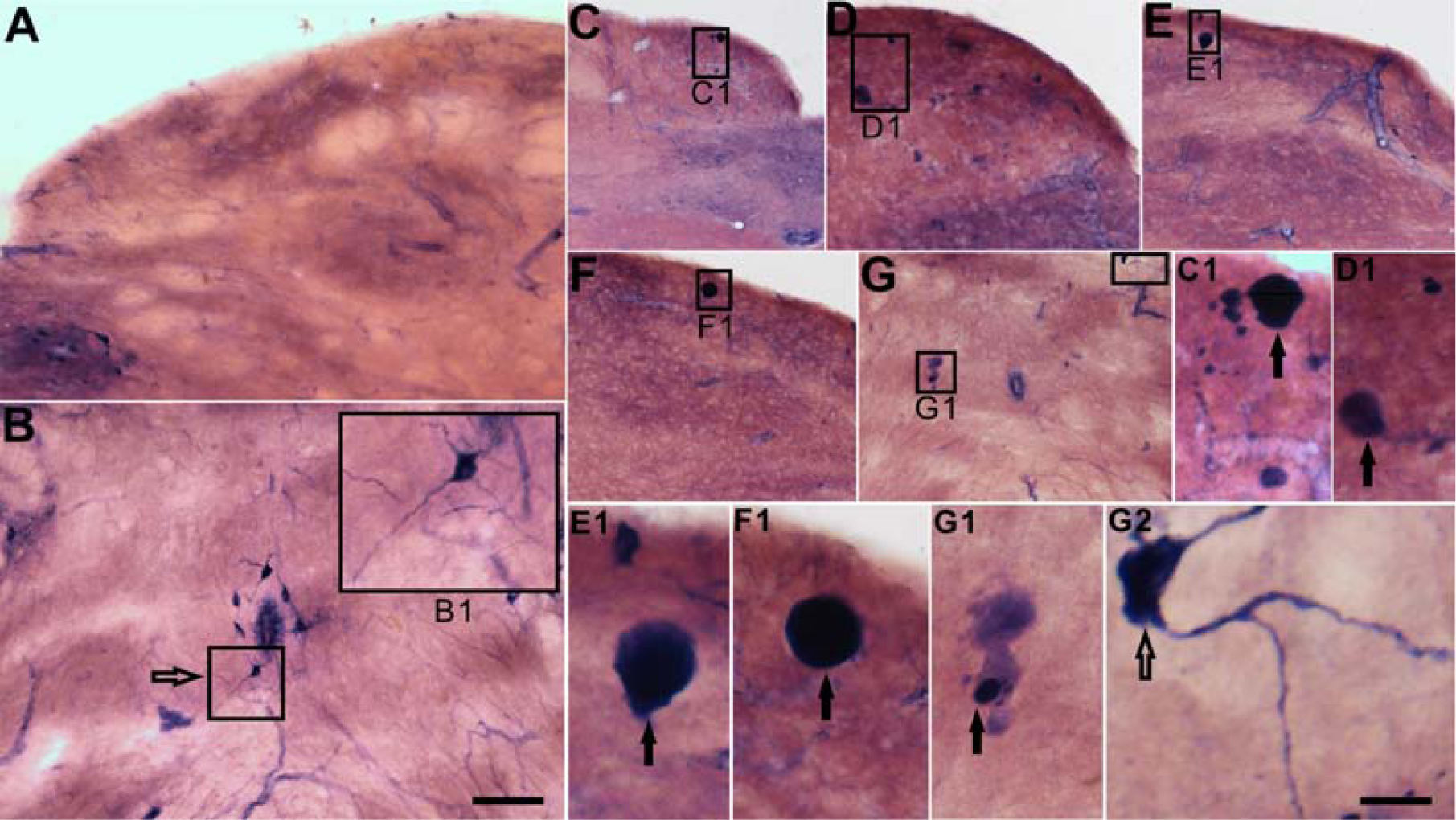
Neurogenerative N-d reactivity in aged (18-month-old) and young adult (2-month-old) mice. (A, B) N-d reactivity in nucleus gracilis and the sacral cord of dorsal commissural nucleus in young mouse. Open arrows indicate N-d positive neurons at nucleus gracilis and the sacral spinal cord of young animals. (C-E) ANB (black arrow) at nucleus gracilis of aged mouse. (C1-F1) refers to the different forms of ANB in the nucleus gracilis. ANB in G1 and N-d positive neurons of G2 in the sacral cord of dorsal commissural nucleus of aged mouse. Sacral bar=50μm.

In order to further study of immunohistochemical profiling of the spinal cord in the young and aged pigeons, NeuN, GFAP, Iba1, MAP2, VIP and CGRP were further examined by immunofluorescence (Figure 10, 11). Omitting of the primary antibodies, all control tissue sections exhibited no evidence of immunoreaction for control of immunofluoresent staining. Mouse anti-CGRP was failed to visualize any immunoreaction (data not showed here), which may be the species-specific reason. Compared with adult animals, the expression of VIP, GFAP and Iba1 were increased in the lumbosacral spinal cord of aged pigeons while the expression of NeuN and MAP2 were decreased (*p* < 0.05). The same result was found in the other spinal segments (data not showed here).

**Figure 10.**
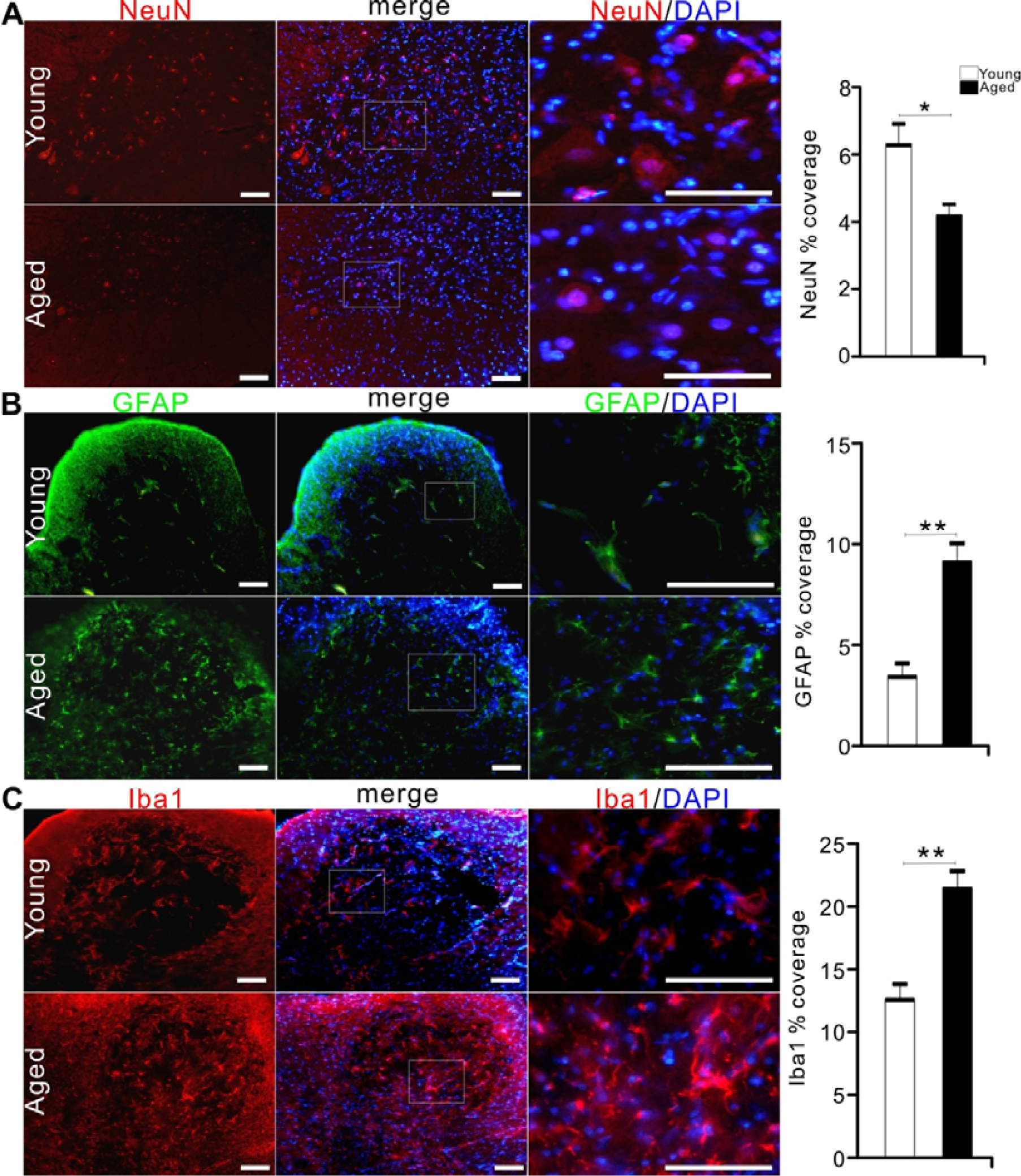
Immunofluorescent profile in the dorsal and ventral horn of lumbosacral segments of young and aged pigeons. Y-axis in bar plot shows percentage coverage of immunoreactive area within the selected area for NeuN, GFAP, Iba1. (A) Fluorescent immunostainig for NeuN and DAPI of lumbar ventral horn. (B) Fluorescent immunostaining for GFAP and DAPI of the sacral dorsal horn. (C) Fluorescent immunostaining for Iba1 and DAPI of lumbar dorsal horn. \**p* < 0.05, \*\**p* < 0.01. Scale bars=50μm.

**Figure 11.**
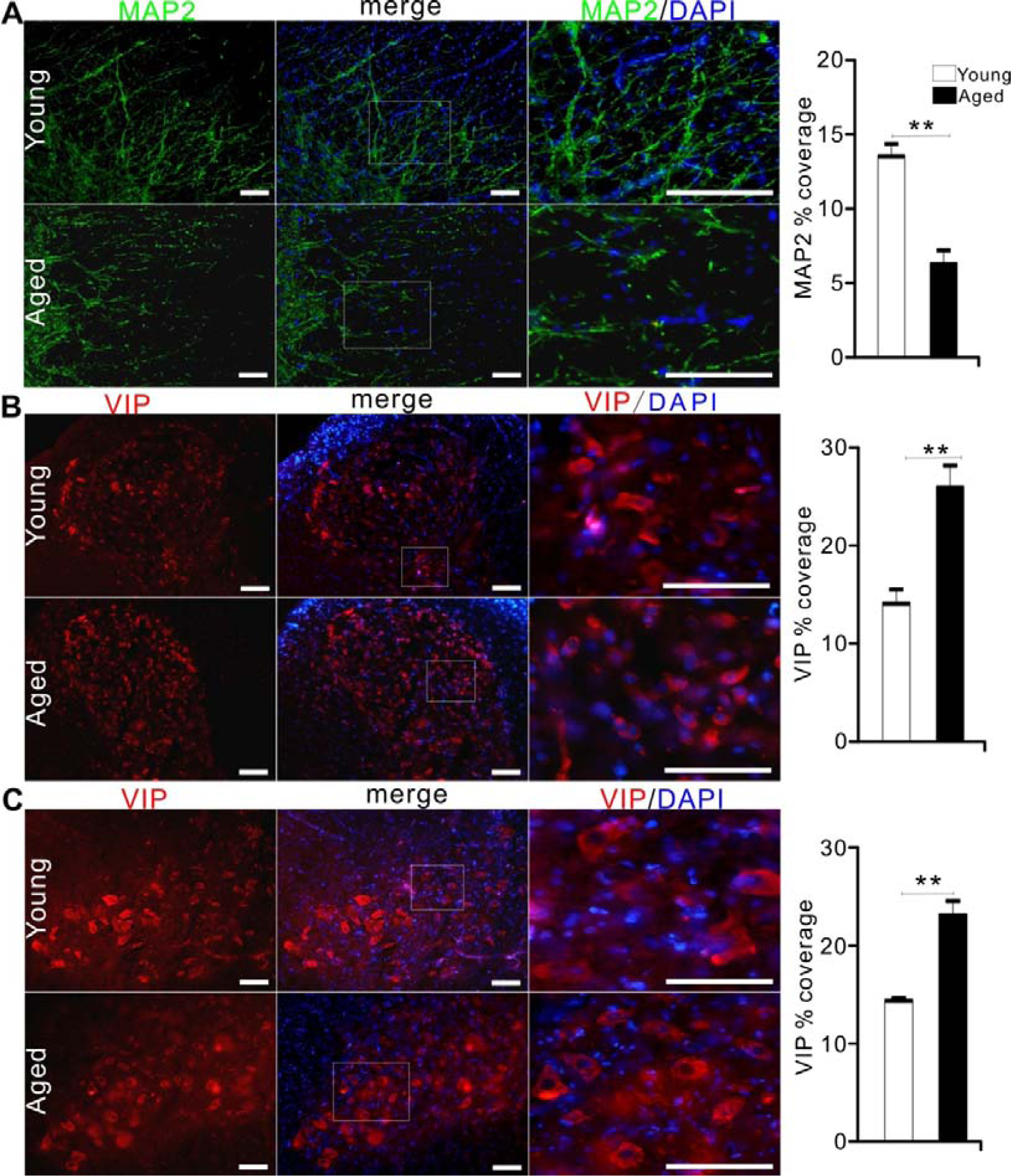
Immunofluorescent profile in the dorsal and ventral horn of lumbar segments of young and aged pigeons. Y-axis in bar plot shows percentage coverage of immunoreactive area within the selected area for MAP2 and VIP. (A) Fluorescent immunostaining for MAP2 and DAPI in the funiculus lateralis of the lumbar segment. (B) Fluorescent immunostaining for VIP and DAPI in the dorsal horn of lumbar segment. (C) Fluorescent immunostaining for VIP and DAPI in the ventral horn of lumbar segment. \**p* < 0.05, \*\**p* < 0.01. Scale bars=50μm.

## DISCUSSION

N-d neurons play important in the somatic and visceral sensation (Valtschanoff et al., 1992; Maisky et al., 2002; Ma et al., 2016). Pigeon behavioral responses showed higher threshold in von Frey test and insensitive to nociceptive hotplate testing in our experiments. This result indicated that the somatic sensory system of pigeons was different from other species. To our knowledge, this is the first time to study the cellular types and quantification of N-d positivity combined with other important neurochemistry in the spinal cord and medulla oblongata of young and aged pigeon. The lumbosacral specialization related to the central canal and related architecture is also described confirmedly with N-d histology and Nissl staining. N-d neurons in anterior commissure are identified which may play an important role in information exchange on both sides of the gray matter. The optimistic N-d staining reveals a Golgi impregnation-like quality (Valtschanoff et al., 1992). Actually, most of the N-d neurons in the spinal cord and dorsal column nuclei of pigeons did not define the positive neurons with a Golgi impregnation-like manner in our experiment. Many of N-d neurons occurred dense or moderate stained soma were barely visible processes. The increased N-d positive neurons in the lumbosacral of white matter relative to the cervical、thoracic region suggest that it may be involved in the visceral regulation of pigeon. ANB in the CuE suggested neurodegeneration in the nervous system of aged pigeon. At least, it can be demonstrated that ANB was prevalently among different species. ANB as an aging marker is identical to the avian and mammals.

This study shows that N-d positive cells in young animals is consistent with previous studies (Necker, 2004). Cutaneous nerves and feather follicles were represented predominantly in laminae □ and □ (Wild, 1985; Woodbury, 1992). Laminae □-□ of the pigeon received somatosensory primary afferents from the skin and muscles of the leg or wing (Necker, 1985; Wild, 1989). Through the analysis of statistical data, we know that N-d cells are mainly distributed in the dorsal horn, especially in the lumbar segments. The lumbar spinal cord not only participates in the regulation of proprioception, but also in the regulation of pelvic organs. For cell body size, it is the largest in the lumbar ventral horn, which may be due to the complex motor function, continuous sensory feedback and serves to integrate of the spinal cord circuits (Atoji et al., 1996). For aged pigeons, the number of cells decreased or the size of cell body reduced. It was postulated that some functions of the spinal cord were deterioration. We divided the N-d positive cells into two types according to the number of dendrites, type □ and type □, which is quite different from the mammal those with rich dendrites (Valtschanoff et al., 1992). But the pigeon has its own particularity, that is the so-called ‘‘Terni’s column’’ (Macchi et al., 2013). It was founded by Terni and named it. He described the longitudinal column of nervous cells between the first thoracic and the second lumbar segments of the spinal cord of a preganglionic nervous center, which receive visceral afferent inputs and innervate visceral organs (Cabot and Cohen, 1977; Ohmori et al., 1987). In this study, the N-d in the CT was positive, but cell morphology is different from intermediolateral column (IML) in mammals with lavish dendrites (Valtschanoff et al., 1992). Few studies have revealed the characteristics of CT from N-d cellular morphology and explored its possible functions. We also found that there was a magnocellular column in the fifth lamina of the enlargement spinal cord, which could also be called CIC. CIC is a major relay center for unconscious proprioception (Necker, 1990a). Sensory information for muscle spindles and tendon, organs, synapses onto posterior root ganglion, which in turn synapses onto Clarke’s column (Necker, 1990b). The cells in Clarke’s column have been shown to be the cells of origin of spinocerebellar pathways (Oscarsson, 1973). The perikaryon area is indicated as the largest of all N-d positive cells. This may be consistent with the ability of birds to maintain balance in flight.

With regard to the spinal cord of a bird, a specialization of pigeon spinal cord is semicircular canal-like structures in the coronal transverse section through the glycogen body (Matulionis, 1972; Necker, 1999; Moller and Kummer, 2003; Imagawa et al., 2006). Research suggests the correlation of function of the blood-brain barrier with the glycogen body (Moller and Kummer, 2003). In addition, other studies suggest that the lumbosacral specialization of the vertebral canal with the spinal cord as a whole system is thought to function as balance sensation (Necker, 1999). But few people have revealed it from the results of N-d staining in the lumbosacral enlargement. There is almost no data to explain how the diamond groove is open and how to close. This paper may give some explanations from the perspective of morphology. First, a cell-free band was formed above the central canal, then the strip was gradually widened and split and filled with the appearance of the colloidal substance. Meanwhile, the dorsal cord on both sides is still connected. With the increase of colloidal substance, the gray matter and dorsal cord on both sides are completely separated caudal to the lower end of sacral spinal segment. Splitting the sacral nervous parenchyma gradually merged at the caudal spinal cord.

Our hypothesis pointed out that N-d alterations in the gracile nucleus (Ma et al., 1997) was correlated to ANB in the sacral segment (Tan et al., 2006) of the aging rats. We also found another kind of N-d neurodegeneration, megaloneurites (N-d meganeurites) in aged dogs (Li et al., 2018). Different to ANB in aged rats, N-d meganeurites are swelling and larger diameter neurites occurred in the dorsal root entry zoom, dorsal horn, lateral collateral pathway, Lissauer’s tract, IML and DCN in aged dogs (Li et al., 2018). In addition, ANB occurred in the GC and CuE of aged pigeon, but not in the lumbosacral segment. In general, innervation of the lower extremities is somatotopically confined to the GR, whereas that of the wing already extends laterally, from the CU into the CuE, which also receives somatosensory fibers and upper half of the body input from the cutaneous and muscular sources in the pigeon. Reasonably, the CU of pigeons is much larger than that in rats. Somatosensory system damaged in elderly pigeons due to various harmful factors. This neurodegeneration can be well demonstrated by N-d histochemical staining, which is bound to be accompanied by a decline in the function of normal N-d positive neurons. The decrease in the number of normal N-d positive neurons in the CuE is also corroborated with age. The aged pigeons in this study may be not old enough, or pigeons have a strong ability to resist oxidative stress, which caused that the number of ANB was relatively less than that of rats (Coppola et al., 2014). In addition, it may be related to insensitivity of pigeon sensory system. Our behavioral tests of pigeons (data not showed here) could be considered the presumption. As far as the lumbosacral spinal cord of aged rats is concerned, the location of ANB is related to the regulation of pelvic organs. The special structure of the lumbosacral spinal cord and the abundant circulation of cerebrospinal fluid and the loss of sacral parasympathetic nucleus in pigeons may explain the absence of ANB in the lumbosacral segment of pigeons. Meanwhile, the pattern of aging neuronal changes in the avian brain is different from that observed in the mammalian brain (Coppola et al., 2016). For the split of the dorsal commissural gray, the N-d positivity of the medulla and lumbosacral spinal cord in birds showed the aging discrepancy from that of mammals. Or, referred to a smaller number of ANBs in the GC, it was also considered for insensitive to thermal and mechanical sensation in the aged pigeons.

In summary, two noticeable subgroups of N-d cells were identified: type □ neuron and type □ neuron. The major new finding revealed by this study was that N-d neurons were enormously distributed in lamina □, and lamina □ and □ and scattered neurons in the ventral horn as well as the white matter of every spinal segment. Moreover, the N-d neuron pool was mostly distributed in the lumbar segments. Referring to aged pigeon, the number of N-d neurons was reduced in most spinal segments. Aging changes also occurred in cell types and immuncytochemistry. The avian neuronal aging deterioration of N-d activity in the lumbosacral spinal cord was different from that of the mammals, but ANB was identified in the medulla oblongata of both aged murine and aged pigeons. It is speculated that the alteration of N-d histology is considered as a marker of neurodegeneration in diverse species.

## Supporting information

Figure S1. Regional indication for example for N-d staining of the pigeon spinal cord.

## ACKNOWLEDGMENTS

This work was supported by grants from National Natural Science Foundation of China (81471286), Undergraduate Training Programs for Innovation and Entrepreneurship of Liaoning (201410160007) and Research Start-Up Grant for New Science Faculty of Jinzhou Medical University (173514017). The authors have no conflicts of interest to declare.

