## Supplementary figures and images for "A comparative assessment of aging-related NADPH diaphorase positivity in the spinal cord and medullary oblongata between pigeon and murine"

### Figure S1. Regional indication for example for N-d staining of the pigeon spinal cord.

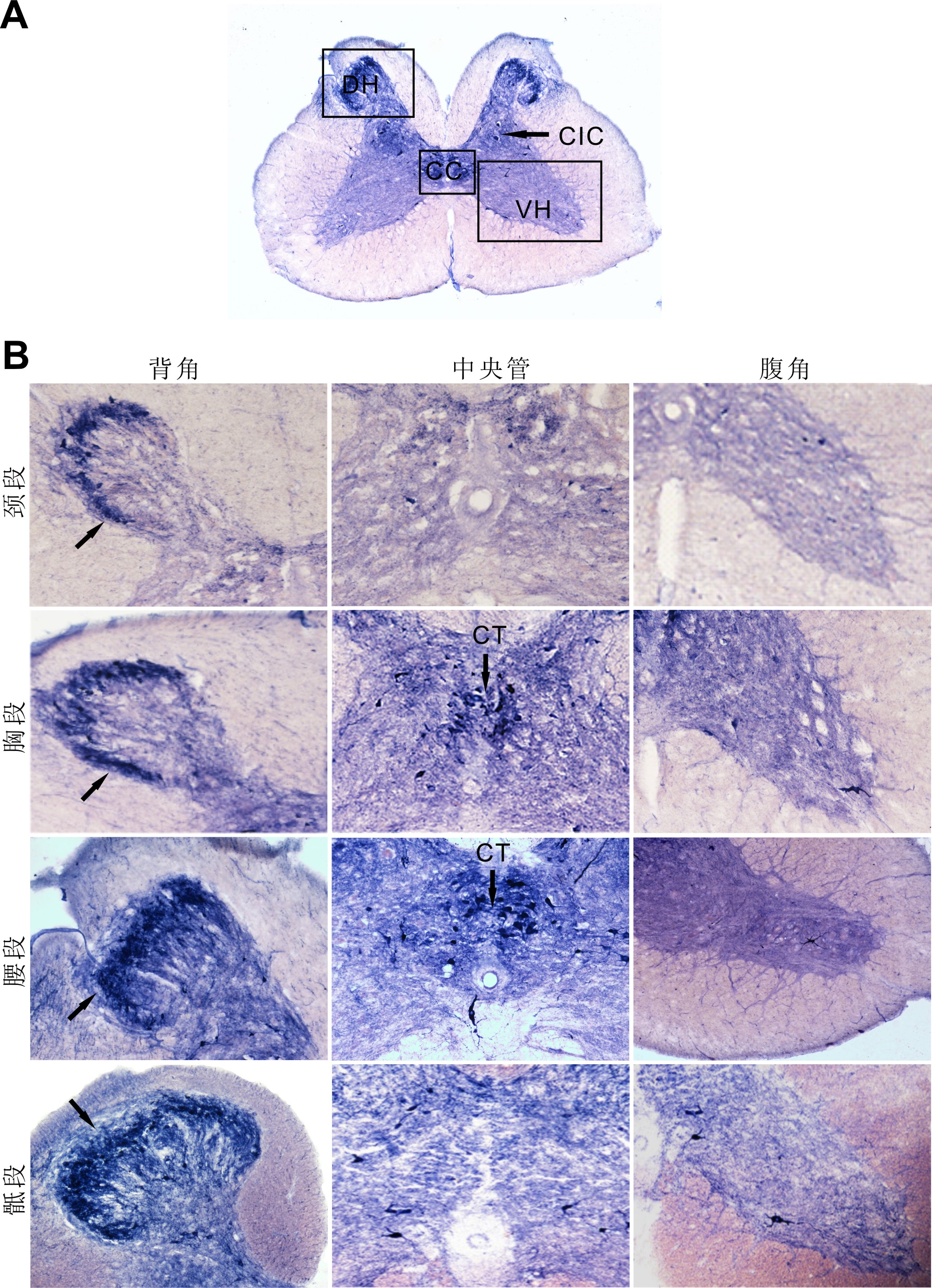
